# Genotoxin-induced transcriptional repression regulates selective protein aggregation

**DOI:** 10.1101/116822

**Authors:** Veena Mathew, Annie S. Tam, Karissa L. Milbury, Analise K. Hofmann, Christopher S. Hughes, Gregg B. Morin, Christopher J. R. Loewen, Peter C. Stirling

## Abstract

Upon genotoxic stress, dynamic relocalization events control DNA repair, and alterations of the transcriptome and proteome enabling stress recovery. How these events may influence one another is only partly known. Beginning with a cytological screen for genome maintenance proteins that move under stress, we find that, upon alkylation stress, the splicing factor Hsh155 localizes to both intranuclear and cytoplasmic protein quality control aggregates. Under stress, an ordered sequestration of Hsh155 occurs at nuclear and then cytoplasmic aggregates in a manner that is regulated by molecular chaperones. This dynamic behavior is preceded by a decrease in splicing efficiency. While DNA replication stress signaling is not required for Hsh155 sequestration, Hsh155 aggregation is cell cycle and TOR pathway dependent. Indeed, loss of a TORC1 regulated ribosomal protein gene transcription factor Sfp1 allows general aggregate formation but prevents Hsh155 recruitment. Together, our analyses suggest a model in which some proteins evicted from chromatin undergoing transcriptional remodeling during stress are targeted to protein quality control sites.

## Introduction

In order to survive chemical or environmental challenges, organisms have evolved a robust network of stress responses that remodel the transcriptome and proteome to enable recovery and, in some cases, future tolerance (Guan et al, 2012). While damaged proteins and RNAs can be turned over, damaged DNA must be repaired in order for cells to continue dividing. However, these DNA repair reactions take place in the context of global changes to the transcriptome and translatome, which help to arrest the cell cycle, and promote survival (Begley et al, 2007; Gasch et al, 2001; Tkach et al, 2012). In *Saccharomyces cerevisiae* (budding yeast), one of the hallmarks of transcriptome remodeling under stress is the suppression of ribosome production which occurs under various conditions and is regulated by TOR-dependant modulation of transcription factors such as Sfp1 (Gasch et al, 2000; Jorgensen et al, 2004; Marion et al, 2004). The ways in which transcriptome alterations impact proteome dynamics under any given stress condition are only partly understood.

Recent genome-wide cytological screens of yeast strains expressing GFP fusions to most proteins have revealed that genotoxin-induced protein relocalization events effect hundreds of proteins, many of which are not directly involved in DNA repair (Chong et al, 2015; Mazumder et al, 2013; Tkach et al, 2012). For instance, in addition to canonical DNA repair foci, treatment of yeast cells with methyl methanesulfonate (MMS) induced novel peri-nuclear foci containing the proteins Cmr1, Hos2, Apj1 and Pph21 (Tkach et al, 2012). These foci were subsequently recognized to be sites of molecular-chaperone regulated protein quality control (PQC) and renamed intranuclear quality control sites (INQ) (Gallina et al, 2015; Miller et al, 2015). Dozens of proteins have now been localized to the INQ, some of which, such as Mrc1, play a role in recovery from genotoxic stress. Subsequent work on Cmr1 and Hos2 found evidence of roles for these proteins in regulating global transcription in unstressed cells, but failed to make connections of their transcriptional roles to their localization at INQ (Jones et al, 2016). The localization and composition of PQC sites in yeast depend upon many factors including the specific type of stress, growth phase, cellular age, and the substrate under study (Saarikangas & Barral, 2016). Studies of both endogenous proteins and aggregation prone model substrates in actively growing cells have identified at least three classes of yeast PQC compartments: INQ/JUNQ (juxtanuclear quality control); CytoQ (cytoplasmic quality control) and the insoluble protein deposit (IPOD). The key players mediating creation and dissolution of these structures are molecular chaperones, in particular small heat shock proteins such as Hsp42 which promote aggregation, and the disaggregation machinery, including Hsp104 and Hsp70 family members (Reviewed in (Saarikangas & Barral, 2016)).

Beginning with a focused screen of >600 yeast chromosome instability proteins, here we uncover INQ localization of a core-splicing factor, Hsh155, upon MMS treatment. We establish the dynamics of Hsh155 sequestration and define a regulatory network of proteins controlling its relocalization. Furthermore, we link Hsh155 aggregation to transcriptional repression of ribosomal protein (RP) genes in MMS conditions, which dramatically alter the need for splicing in yeast. These observations suggest unappreciated influence of transcriptome changes on the composition of PQC sites under stress. We propose a model where repression of RP genes, and the concomitant drop in the need for splicing, liberates Hsh155 and other factors for sequestration at PQC sites in a TORC1 dependent manner. Together our data provide new links between transcriptome regulators and PQC site composition under stress, where stress activated TORC1 regulated transcriptional program is controlling the composition of PQC sites.

## Results

### A screen for genome stability factors that relocalize after DNA damage identifies Hsh155

To explore dynamic responses of proteins with known roles in genome integrity to genotoxic stresses, we screened a biased mini-array of GFP-fusion proteins comprised entirely of proteins whose mutation is linked to an increase in genome instability (Stirling et al, 2011; Stirling et al, 2014). The 632 GFP-tagged genome maintenance proteins were imaged at high resolution after no treatment or exposure to the alkylating agent MMS, ultraviolet (UV) irradiation, or H_2_O_2_. Candidate relocalization behaviors from the primary screen were validated in triplicate leading to a final list of 41 relocalization events after genotoxic stress (**Supplementary Table S1**). Most relocalization events occurred in all three stresses, and a large majority occurred in at least two conditions, with only 8 appearing under a single stress condition (**Fig.1A**). Comparison of our data to three previously published genome-wide MMS and H2O2-induced relocalization screens show a high degree of overlap (i.e. 5/41, 19/41 and 10/41 respectively) (**Fig. 1B**) (Breker et al, 2013; Mazumder et al, 2013; Tkach et al, 2012). Most movements occur into or out of the nucleus, or into nuclear or cytoplasmic foci (**Fig.1C, Supplementary Fig. S1**), which, based on their annotation (www.yeastgenome.org), we can ascribe to the formation of aggregates, P-bodies, or DNA repair centers. One unexpected observation was the relocalization of Hsh155-GFP into nuclear and cytoplasmic foci in response to MMS or H_2_O_2_ (**Fig.1D and E**). Hsh155 is part of the SF3B subcomplex in the U2-small nuclear ribonucleoprotein (snRNP) of the spliceosome. To assess the specificity of Hsh155 relocalization, we tested the relocalization behavior of Hsh155 binding partners within the spliceosome, Cus1 and Hsh49, after MMS treatment but did not observe any change in their localization (**Fig.1F**). Nuclear Hsh155 foci did not colocalize with canonical DNA damage repair proteins making a direct role in DNA repair unlikely (**Supplementary Fig. S2A**). Our data thus defines a previously unrecognized and selective relocalization of Hsh155 to foci after alkylating and oxidative genotoxic stress.

**Figure 1.**
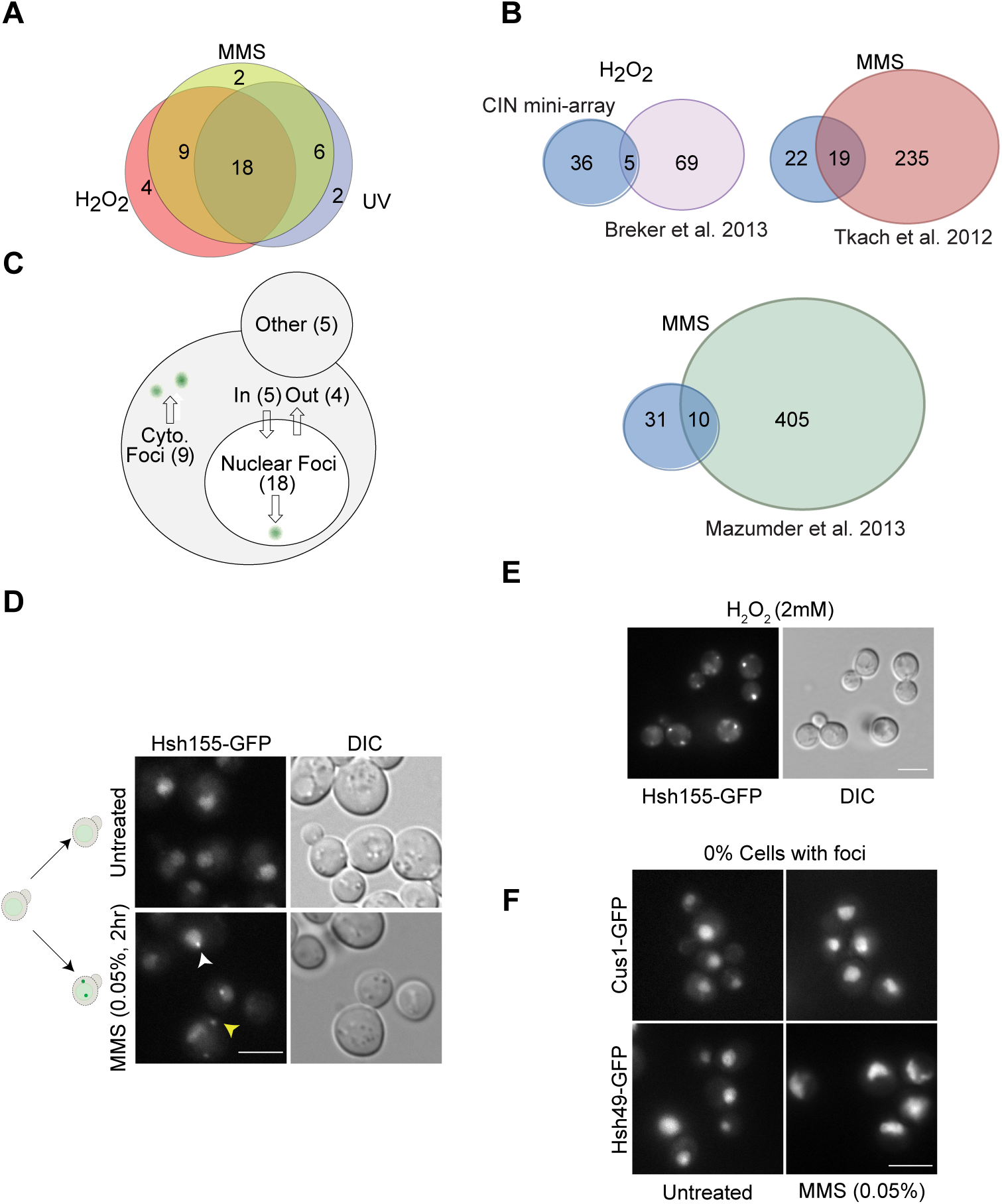
DNA damage relocalization screen of the chromosome instability (CIN) proteome identifies Hsh155. (A) Overall screen results. A Venn diagram of overlapping protein relocalization behaviors upon MMS, H_2_O_2_ or UV treatment. List of proteins relocalized in each treatment detailed in **Supplementary Table S1**. (B) Comparison of screen results with published whole proteome relocalization screens. The stress is indicated above and the reference below. (C) Yeast cell schematic summarizing relocalizations by destination under stress (see also **Supplementary Fig. S1** for sample images of novel localization, and **Supplementary Table S1**). (D) MMS induced relocalization of Hsh155 into nuclear (white arrowhead) and cytoplasmic foci (yellow arrowhead). A schematic (left) summarizes the movements. (E) H_2_O_2_ induced relocalization of Hsh155. (F) U2-snRNP spliceosome complex partners of Hsh155, Cus1 and Hsh49 do not form foci after MMS treatment. Scale bar is 5μm for all subsequent figures.

### Hsh155 foci are protein aggregates

The formation of a single focus peripheral to the nucleus by Hsh155 was similar to the recently described intranuclear quality control (INQ) compartment, which contains Cmr1, Hos2 and many other proteins (Gallina et al, 2015; Miller et al, 2015; Tkach et al, 2012). Indeed, Hsh155-GFP colocalized with Hos2-mCherry at both cytoplasmic and nuclear foci after MMS (**Fig.2A**). To confirm that these were indeed sites of protein aggregation, we also co-localized Hsh155-GFP with VHL-mCherry, a protein that cannot fold in yeast and is targeted to aggregates under various stresses (McClellan et al, 2005). Hsh155 and VHL were significantly co-localized in MMS (**Fig.2B**), although Hsh155-GFP did not join VHL foci at high temperature alone (**Supplementary Fig. S2B**). The aggregates of both Hos2 and VHL have been shown clearly to localize in both nucleus and cytoplasm (Miller et al, 2015). To differentiate and confirm the compartmental sequestration of Hsh155, we further used Hta2 (histone H2A) and Nic96 (nuclear pore protein) as markers of nuclear area (**Supplementary Fig. S2C).** Consistent with localizing in an aggregated state, fluorescence recovery after photobleaching (FRAP) analysis of nuclear Hsh155-GFP or Hos2-GFP foci confirmed a large immobile fraction for each protein (~ 50%) similar to known aggregates of PQC (Saarikangas & Barral, 2015), and a recovery time (t_1/2_ ~25s) much slower than freely diffusing proteins (**Fig. 2C**). These results thus identify the core-splicing factor Hsh155 as a new constituent of the INQ protein aggregate compartment following genotoxic stress.

**Figure 2.**
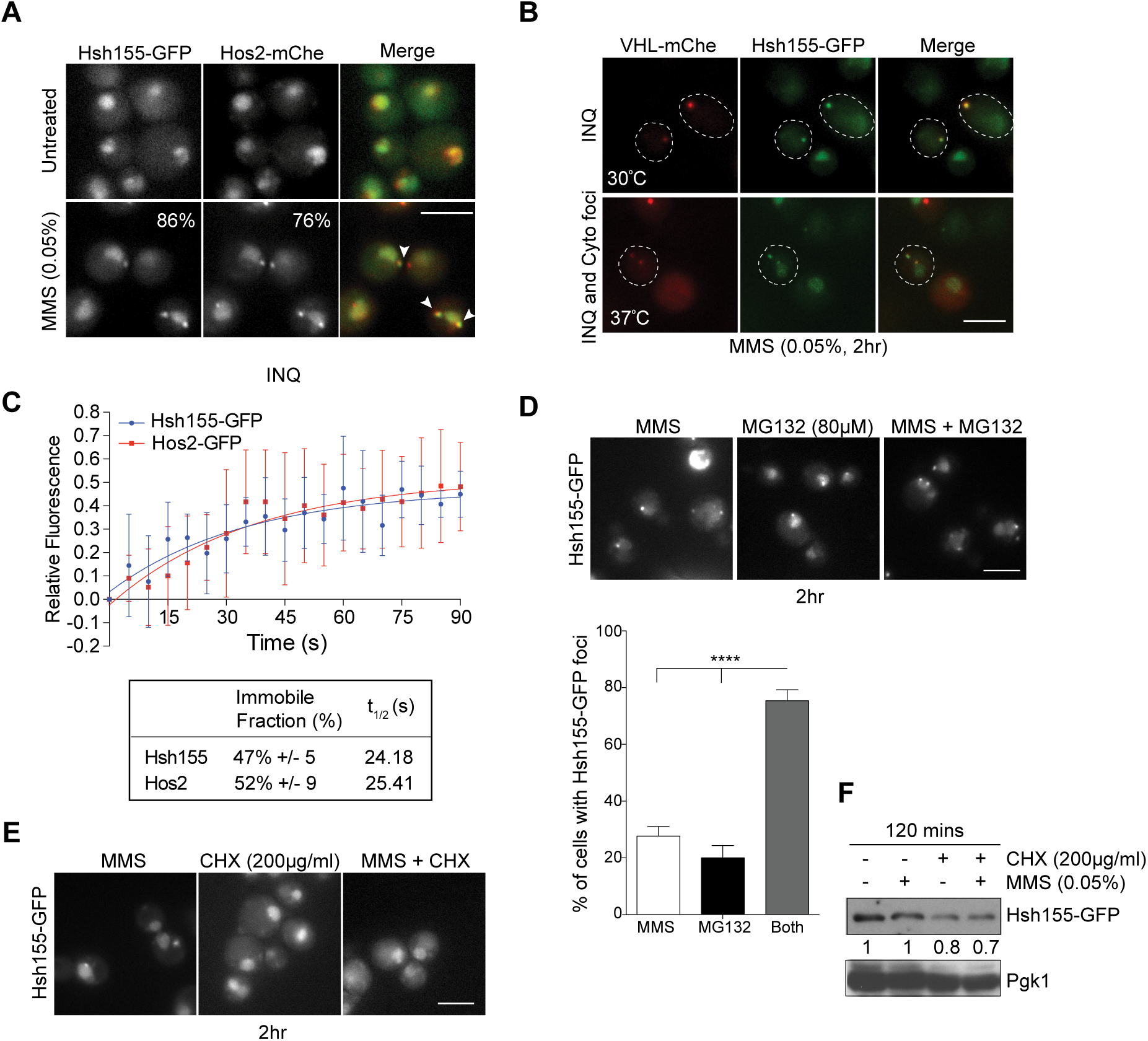
Hsh155 relocalizes to nuclear (INQ) and cytoplasmic protein quality control (PQC) sites. (A) Co-localization (white arrowheads in merge) of Hsh155-GFP with Hos2-mCherry (mChe) in MMS. Inlets show percentage overlap of foci in each to one another. (B) Co-localization of Hsh155-GFP with VHL-mCherry at both 30°C and 37°C with MMS. (C) Quantitative FRAP analysis of GFP tagged Hsh155 or Hos2 in nuclear foci. Top: The best line of fit curve of relative fluorescence intensities over time for both Hsh155 (blue) and Hos2 (red); bottom: percentage of Hsh155 and Hos2 in the immobile fraction and diffusion time (t_1/2_). Values are mean± SD of Hsh155 (8 cells) and Hos2 (5 cells) analysed over three independent experiments. (D) Effect of proteasome inhibition by MG132, inducing Hsh155 aggregation at INQ. MG132 treatment induced foci and is quantified below with or without MMS addition. Shown are the mean values from three independent experiments ± SEM with >100 cells each. (Asterisks show *p*-value (****) p<0.0001, Fisher test). (E) Cycloheximide blocks Hsh155 foci formation. Representative image of Hsh155-GFP tagged cells treated with translation inhibitor, cycloheximide (CHX, 200μg/mL) and MMS (0.05%) with no foci formation compared to MMS treatment alone. (F) Hsh155 protein levels by anti-GFP western blotting relative to Pgk1 levels (as indicated) in cells treated with MMS and/or CHX. Shown is the representative blot from at least three independent experiments.

To examine the effects of new protein synthesis or degradation on the Hsh155 foci formation, we treated cells with the proteasome inhibitor MG132 or the translation inhibitor cycloheximide (CHX) alone or in combination with MMS. Similar to earlier studies on other INQ components (Gallina et al, 2015), MG132 treatment alone induced Hsh155 sequestration to INQ sites while CHX alone had no effect on Hsh155 localization (**Fig. 2D and E**). Consistently, MG132 in combination with MMS significantly increased the number of cells with foci suggesting that protein degradation works in opposition to aggregate formation or to reduce aggregate lifetime (**Fig. 2D**). Remarkably, combined treatment of cells with MMS and CHX completely abolished Hsh155 foci (**Fig. 2E**). New protein synthesis has been shown to be essential for heat-stress aggregates (Zhou et al, 2014) and it is possible that the same is true for INQ foci aggregation with MMS. In addition, the stability of Hsh155 in cells did not appear to change significantly following MMS treatment suggesting that the influence of MG132 on foci formation may be due to other factors rather than Hsh155 itself being a target of degradation (**Fig. 2F; Supplementary Fig. S2D**).

### Dynamic behaviour of Hsh155 at PQC sites

Stress-induced formation of the INQ compartment was reported relatively recently and its dynamics and relationship with other sites of PQC are not well understood. To explore the dynamics of Hsh155 localization to both nuclear and cytoplasmic PQC sites, we first followed Hsh155 foci formation over time. Hsh155 rapidly accumulates at nuclear foci, followed by gradual increases in the frequency of additional cytoplasmic foci (**Fig. 3A**), while washout of MMS led to a gradual decrease of foci and recovery of normal Hsh155 nuclear localization (**Fig. 3B**). Not only does the frequency of foci increase, but the fluorescence intensities of Hsh155 in nuclear and cytoplasmic foci gradually increased over time until three hours, with cytoplasmic foci starting dimmer at early timepoints but matching INQ intensities by 3 hours (**Fig. 3C; Supplementary Fig. S3A**). These results indicate that the clearance of INQ structures proceeds rapidly after stress removal and that, during prolonged stress, the triage pathway shifts from immediate nuclear deposition to a delayed cytoplasmic aggregate deposition.

**Figure 3.**
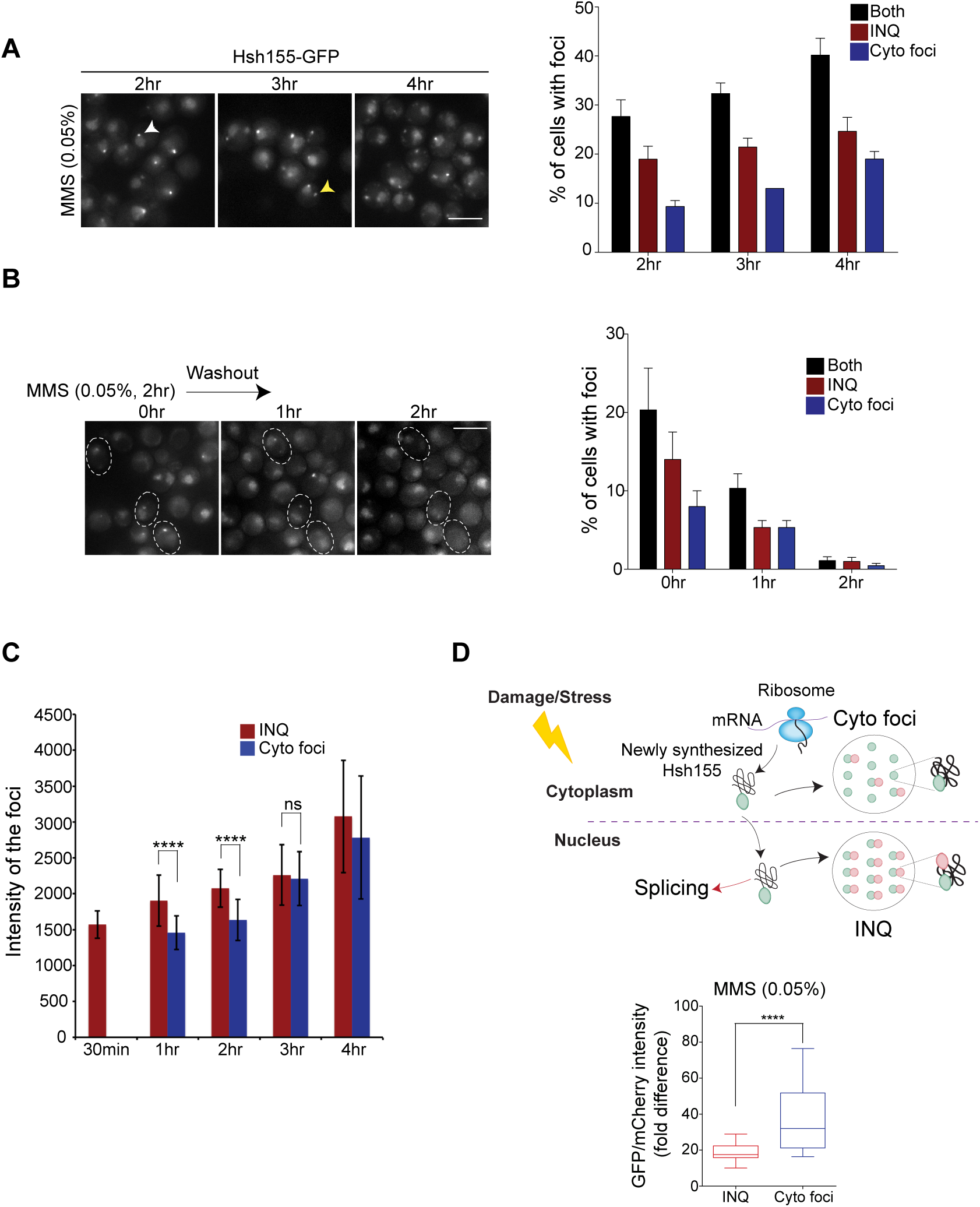
Dynamic behavior of Hsh155 at PQC sites. (A) Time course imaging of Hsh155-GFP foci accumulation shows increase in number of cells with foci over time, predominantly cytoplasmic foci (yellow arrowhead) and (B) disappearance after MMS washout. Representative image are shown (left) and quantification of percentage of cells with Hsh155 foci (right). For (B) dotted inlets represent time-lapse images of foci disappearance over time in the same set of cells. (C) Fluorescence intensity of foci over time show increasing relative GFP in cytoplasmic foci until 2 hr of MMS treatment. (D) Tandem fluorescent fusion intensities of Hsh155 in aggregates. Ratios of GFP (fast maturing) and mCherry (slow maturing) fluorescence in nuclear or cytoplasmic foci are shown (top). A schematic of protein lifetime of Hsh155 fusion to a fluorescent timer in PQC (bottom), showing older (GFP>mCherry) protein in INQ and newer (GFP>>>mCherry) in cytoplasmic foci. Quantification of fluorescent timer indicates aggregation of older proteins in INQ compared to cytoplasm. For A-C, bars are color-coded to denote INQ (red) and cytoplasmic (blue) foci and both (black). For C quantifications: Mean± SEM, three replicates, n> 100, Student t-test, asterisks show *p*-value (****) p<0.0001; (ns) non-significant. For D quantifications: three replicates, n ≥ 28 per replicate, Mann-Whitney test, asterisks show p-value threshold **** p<0.0001.

To assess the aggregation behaviour of Hsh155 over its protein lifetime, we applied a fluorescent reporter tandem fusion approach in which both a fast folding GFP and a slowly maturing mCherry are fused in tandem and the fluorescence ratio of each fluorophore is an indicator of protein turnover rate (Gallina et al, 2015; Khmelinskii et al, 2012). C-terminally tagged Hsh155-GFP: mCherry protein fluorescence ratios in the nuclear or cytoplasmic aggregates were measured after 2 hours in MMS. This experiment revealed that a significantly older pool of Hsh155 (i.e. lower GFP: mCherry ratio) appears in INQ compared to cytoplasmic aggregates (**Fig. 3D**), confirming the sequestration of Hsh155 to INQ first then to cytoplasmic aggregates. The protein turnover rates of the untreated nucleoplasmic signal remain comparable to those after MMS treatment (**Supplementary Fig. S3B**). Together these data reveal the dynamics of Hsh155 under stress and suggest it is first sequestered at INQ sites for refolding and reactivation until stress recovery and only later triaged to cytoplasmic aggregates, possibly more so for nascent Hsh155 since a ‘younger’ pool of protein appears to populate cytoplasmic aggregates.

### INQ resident proteins regulate Hsh155 sequestration

A previous study of INQ aggregates identified several poorly characterized marker proteins that exhibited foci formation under MMS treatment, namely, Cmr1, Hos2, Apj 1 and Pph21 (Tkach et al, 2012). Cmr1 has been implicated in the recovery from DNA damage stress and has been recently linked, together with Hos2, in global transcriptional regulation (Gallina et al, 2015; Jones et al, 2016). Pph21 is one of two protein phosphatase 2A catalytic subunits with pleiotropic functions in the cell, including opposing TOR functions in nutrient signaling (Duvel et al, 2003; Jiang & Broach, 1999). Apj1 is a poorly characterized Hsp40 molecular chaperone family member. To shed light into their functions at INQ, we measured the effects of deleting their encoding genes on Hsh155 foci formation. Deletion of any of these genes significantly increased the number of cells with MMS-induced Hsh155 foci, and shifted the distribution to create more cytoplasmic foci (**Fig. 4A and B**). How each of these INQ markers regulate Hsh155 sequestration may differ and depend on their specific functions in cell (see Discussion). Reciprocally, an *HSH155*-DAmP (Breslow et al, 2008) allele increased the frequency of Hos2-GFP and Apj1-GFP foci in both INQ and cytoplasmic sites after MMS treatment (**Fig. 4C**). Given the core role of Hsh155 in splicing, and the fact that a DAmP allele would simply reduce the amount of wild type protein, suggests that defective splicing may influence the formation of protein aggregates upon MMS treatment.

**Figure 4.**
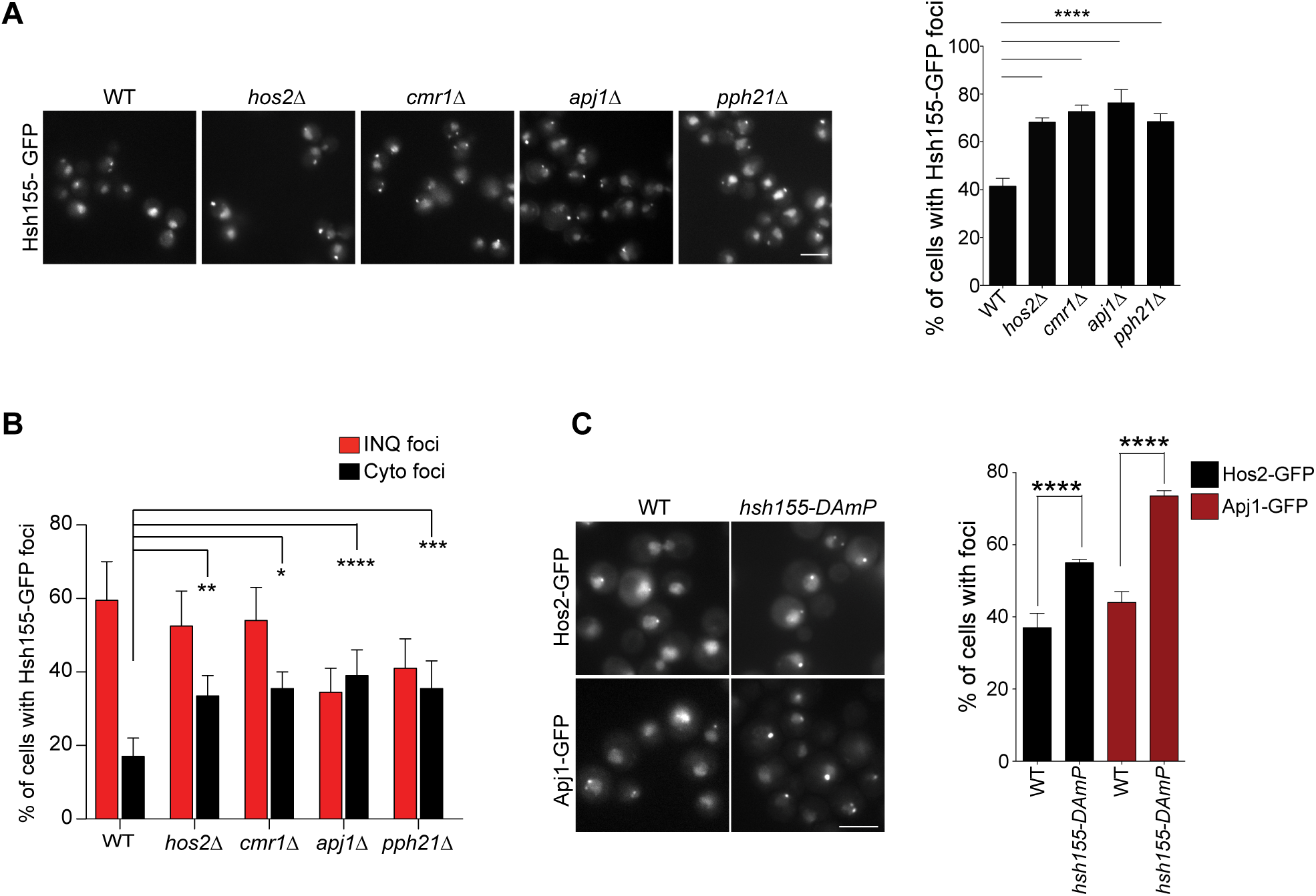
Regulation of Hsh155 foci formation by INQ resident proteins. (A) Representative images showing foci in MMS treated cells in *CMR1, HOS2, APJ1* or *PPH21* deletion strains compared to WT (left). Quantification of percentage of cells with Hsh155 foci in indicated strains (right). (B) Effect of deleting *CMR1, HOS2, APJ1* or *PPH21* on INQ and cytoplasmic foci distribution. Percentage of cells with cytoplasmic foci (black bar) increased significantly in respective mutants compared to WT. (C) Representative images shows effect of depleting *HSH155* on INQ formation (left). Percentage of cells with Hos2 (black) and Apj1 (red) foci in *hsh155-DAmP* or WT cells (right). For A-C quantifications: Mean± SEM, Fisher test, Asterisks show *p*-value thresholds (**** p<0.0001; *** p<0.001; ** p<0.01; * p<0.05).

### A network of molecular chaperones regulates Hsh155 PQC site deposition

Apj1 is a molecular chaperone of the Hsp40/DnaJ family, which our data suggest opposes Hsh155 localization to both nuclear and cytoplasmic foci (**Fig. 4A**). Indeed, chaperones regulate INQ localization of Cmr1 (Gallina et al, 2015). Previous work has implicated compartment specific aggregases Hsp42 and Btn2 in driving substrates to cytoplasmic and nuclear PQC sites respectively (Miller et al, 2015). Surprisingly, deletion of either *HSP42* or *BTN2* almost completely abrogated Hsh155 localization to both INQ and cytoplasmic PQC sites **(Fig. 5A** and **C**). Although the distribution of foci between INQ and cytoplasm in *btn2∆* was similar to WT a slight increase in cytoplasmic foci was seen (**Supplementary Fig. S4A**). While this conflict with reports of compartment specific functions of Hsp42 and Btn2 when analyzing model aggregating substrate proteins (Miller et al, 2015), it is consistent with redundant effects of Hsp42 and Btn2 on Cmr1-YFP foci (Gallina et al, 2015). Therefore, at least for some endogenous protein substrates of INQ, both Hsp42 and Btn2 promote aggregate localization in the nucleus and cytoplasm. Our MMS washout results (**Fig. 3B**) suggest refolding and reactivation of Hsh155 to its native form after stress removal, indicating participation of disaggregases in the process. This led us to interrogate loss of disaggregases Hsp104 and Sse1, which work together in prion destabilization (O'Driscoll et al, 2015). Deletion of *HSP104* and *SSE1* dramatically increased the frequency of PQC sites marked by Hsh155 in MMS, increasing the number of cells with predominantly more cytoplasmic aggregates (**Fig 5B, Supplementary Fig. S4B**). Deletion of *SSE1* led to formation of Hsh155 foci even without MMS (**Fig. 5B**) and shifted the foci to more cytoplasmic aggregates (**Supplementary Fig. S4B**) indicating that Sse1 may be involved in *de novo* folding of Hsh155 or that *sse1∆* yeast experience ongoing stress that affects spliceosome integrity. Together our data show how a network of chaperones acting as aggregases (Hsp42/Btn2) or disaggregases (Apj 1/Hsp104/Sse1) regulate Hsh155 localization to INQ and cytoplasmic PQC sites after stress (**Fig. 5C**).

**Figure 5.**
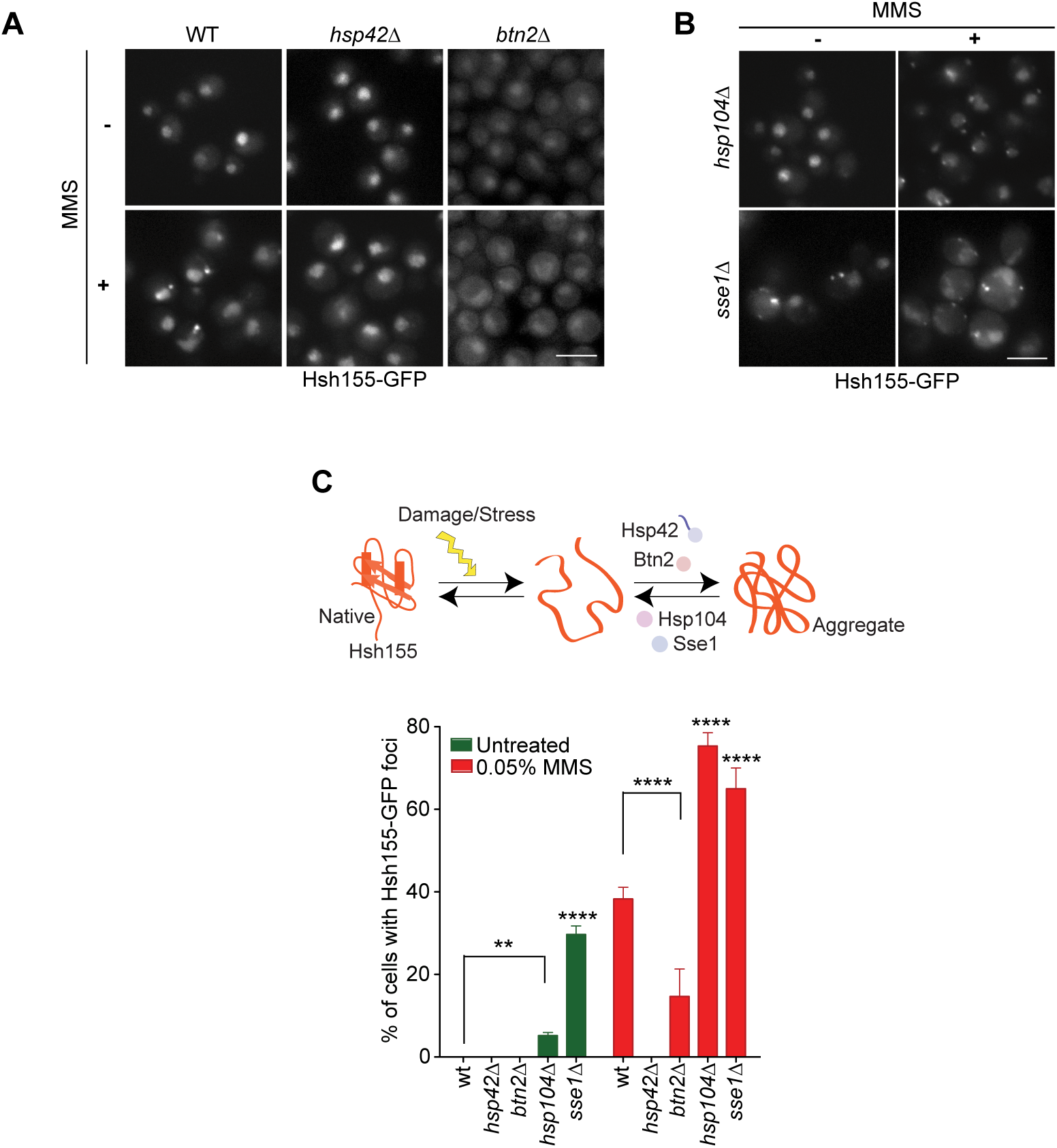
Regulation of Hsh155 foci formation by molecular chaperones. (A) Effect of aggregases Hsp42 and Btn2 and (B) disaggregases Hsp104 and Sse1 on Hsh155 relocalization. Representative images shows decrease (A) and increase (B) in number of cells with foci in the indicated strains relative to WT. (C) Summary of chaperone regulation of Hsh155 relocalization. Schematic (above) and quantification of percentage of cells with foci from A and B (below). Mean± SEM, Fisher test. Asterisks show *p*-value thresholds in comparison to WT under the same condition (** p<0.01; ^*\*\*\*\**^ p<0.0001).

### Hsh155 foci formation is cell cycle dependent

One of the major responses to MMS treatment is DNA replication stress and subsequent signaling propagated through the kinases ATM/Tel1 and ATR/Mec1 (Gasch et al, 2001; Tkach et al, 2012). We wondered whether Hsh155 itself could be required for survival in response to genotoxins, however *hsh155*-temperature sensitive mutants (*hsh155-ts*) did not show a growth defect in MMS, making a direct role in DNA repair unlikely (**Fig. 6A**). Likewise, though we saw an overall increase in signal intensities with hydroxyurea (HU), which induces replication stress without the chemical DNA lesions induced by MMS, consistent with a recent report (Chong et al, 2015), we failed to observe any significant foci formation, suggesting that stalling DNA replication alone is insufficient for Hsh155 foci formation (**Fig. 6B**). To interrogate the function of DNA damage signaling in Hsh155 PQC site localization, we measured PQC foci formation in a strain lacking both yeast ATM (*tel1∆*) and ATR (*mec1∆*). Hsh155 foci formed normally in the *mec1∆tel1∆* mutant compared to wildtype or to *sml1∆* (*SML1* deletion permits viability of *mec1∆* strains) cells, suggesting that DNA damage signaling is not required for aggregation (**Fig. 6C**). Interestingly, a significant number of cells exhibited Hos2-GFP aggregates in the *mec1∆tel1∆* mutant even in unstressed cells and this was further enhanced in MMS (**Fig. 6C, Supplementary Fig. S5A**). This shows that Hos2 foci formation also does not require DNA damage signaling but suggests that stress in the *mec1∆tel1∆* mutant is sufficient to promote Hos2 but not Hsh155 aggregation. These data support the idea that while INQ components like Hos2 and Hsh155 can have co-dependent relationships at PQC sites, they can be governed by independent upstream signals driving sequestration to PQC sites. Furthermore, since MMS is known to arrest cells in S-phase (Shirahige et al, 1998), we synchronized cells either in G1 or G2/M prior to MMS treatment to determine whether passage into S-phase was required for foci formation. While α-factor arrested G1 cells formed Hsh155 foci readily upon MMS treatment, nocodazole arrested G2/M cells formed very few foci (**Fig. 6D** and **Supplementary Fig. S5B**). This shows that MMS-induced replication stress is not the trigger for Hsh155 foci since cells need not enter S-phase in MMS to cause Hsh155 foci, consistent with our data showing *mec1∆tel1∆* has no effect on Hsh155 foci. Together these data show that neither MMS-induced DNA replication stress nor canonical DNA damage signaling is necessary for Hsh155 sequestration at PQC sites and the aggregates are likely formed predominately in G1/S cells.

**Figure 6.**
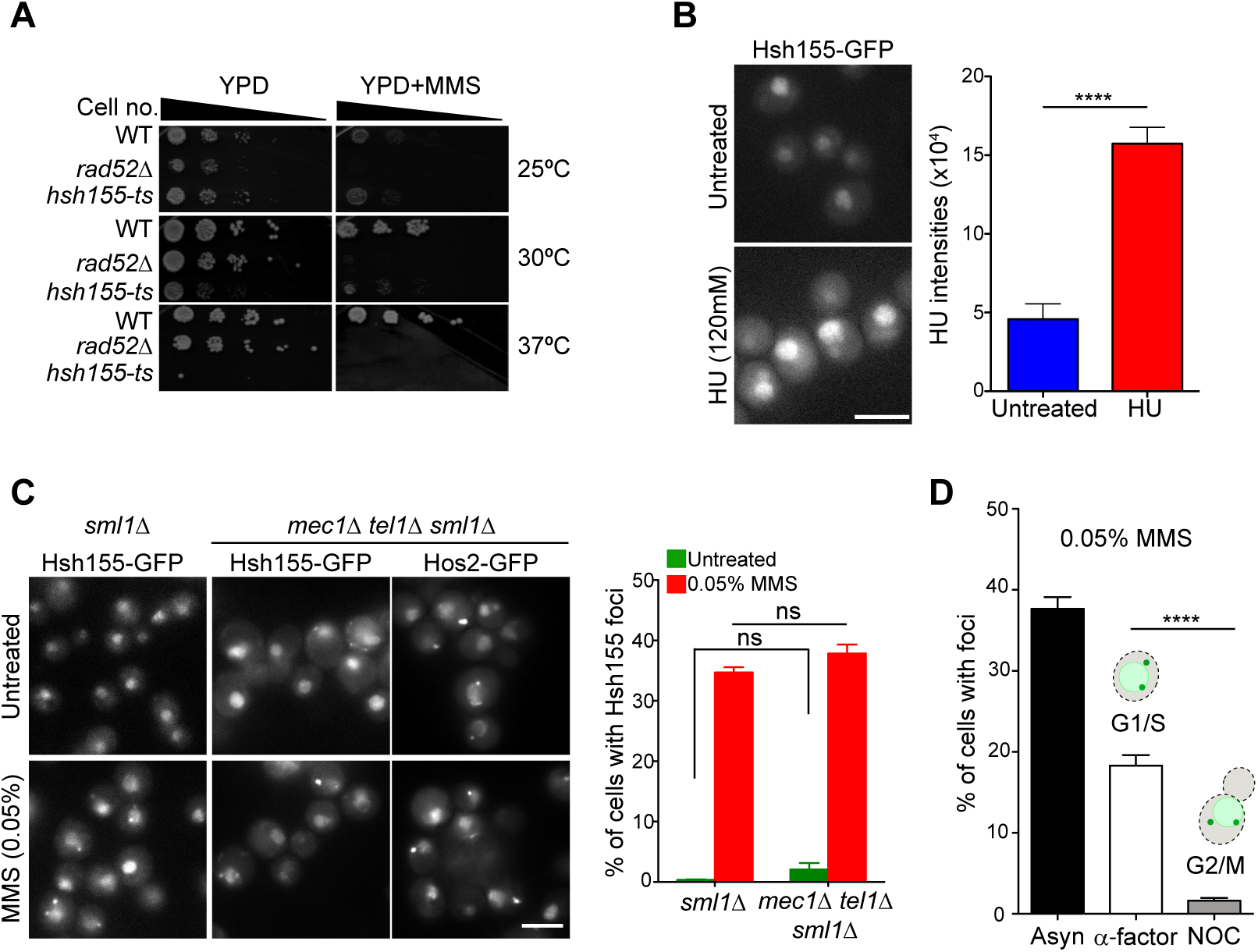
Hsh155 foci are formed predominately in G1 cells. (A) *hsh155-ts* alleles are temperature sensitive but not additionally MMS sensitive compared to WT and *rad52∆* (sensitive control) strains. Equal ODs of the indicated strains were serially diluted and spotted on YPD ± 0.01% MMS at 25°C, 30°C and 37°C. (B) Representative images showing no foci formation but increase in Hsh155 intensities in HU treated (120mM, 2hr treatment) cell compared to untreated. Single cell quantification of mean intensity of Hsh155 protein per cell as depicted in the image. Three replicates, n>100, Mean± SEM, Student t-test, (****) p<0.0001. (C) Loss of *MEC1* and *TEL1* does not influence Hsh155 localization but Hos2 aggregate formation. Bar graph (right) and representative image (left) shows no significant difference in Hsh155 localisation in *mec1∆tel1∆sml1∆* in either untreated (green bar) or MMS treated (red bar) compared to *sml1∆.* Three replicates, n> 100, Mean± SEM, Fisher test, *p*-value (ns) non-significant. (D) Cell cycle dependence of Hsh155 sequestration. Quantification of cells with Hsh155 foci in asynchronous (Asyn, black), G1 (α-factor, white) and G2/M (nocodazole (NOC), grey) cells. Three replicates, n> 100, Mean± SEM, Fisher test, ****p <0.0001.

### Ribosomal protein gene transcriptional regulation and TORC1 influences Hsh155 sequestration

Since Hsh155 is an essential splicing factor we wondered whether the relocalization behavior was linked to the stress-induced remodeling of the transcriptome occurring in MMS. We first analyzed splicing efficiency using a LacZ reporter construct (Palancade et al, 2005) and found that, while MMS treatment reduced LacZ production from both an intronless and intron-containing construct, the intron-containing construct was further repressed, suggesting an additional post-transcriptional splicing defect (**Fig. 7A**). Measuring splicing in strains bearing the *hsh155-*ts allele confirmed that loss of Hsh155 function leads to a dramatic splicing defect (**Supplementary Fig. S6A**). Interestingly, the splicing defect was evident within 30 minutes, well before most cells show detectable INQ foci (**Supplementary Fig. S6B**). This is consistent with the idea that spliceosomes must disassemble in response to MMS treatment prior to Hsh155 aggregation and is supported by observations that other splicing factors tested, including binding partners of Hsh155, did not relocalize in MMS conditions (**Fig. 1F**).

**Figure 7.**
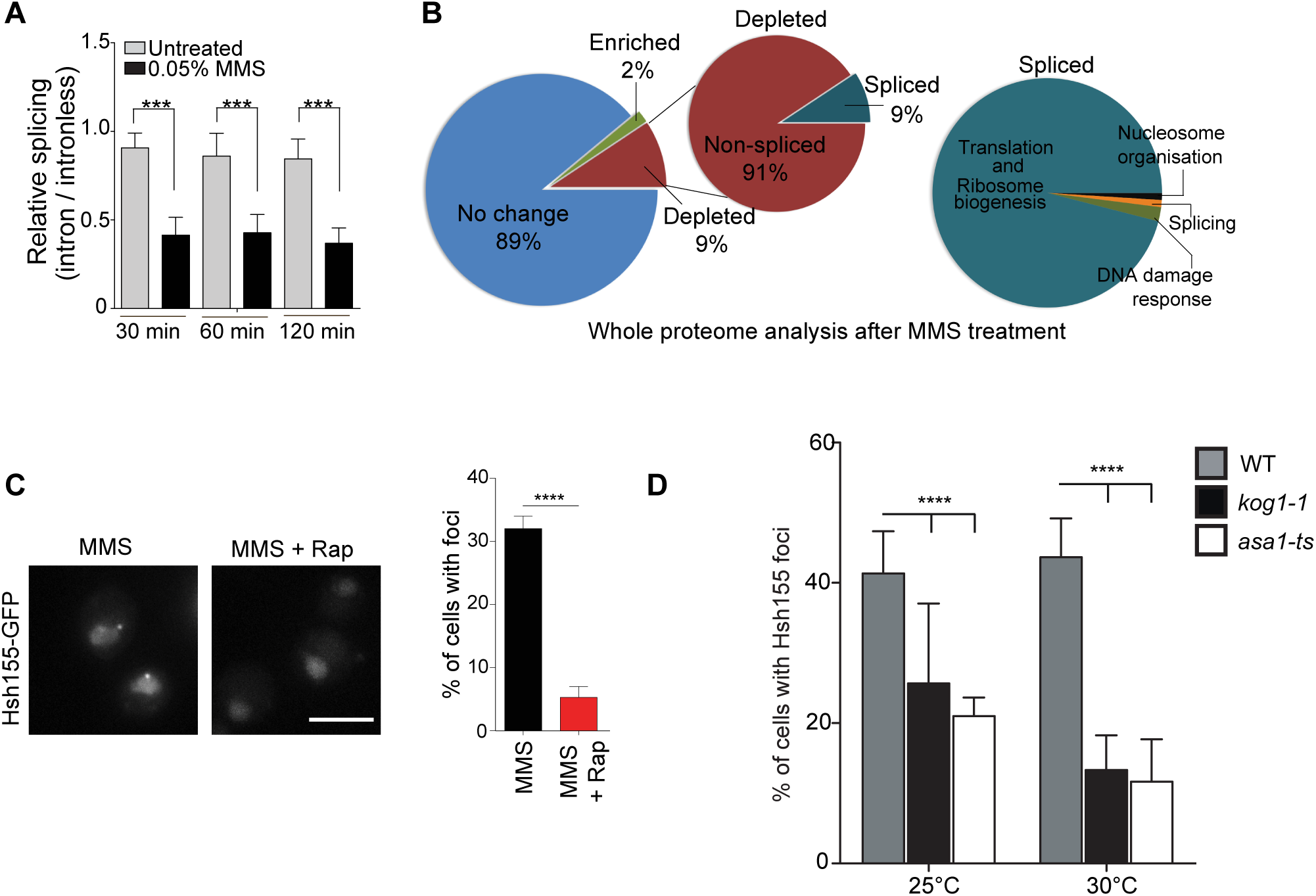
RP gene repression and TORC1 signaling influences Hsh155 sequestration. (A) Splicing efficiency in MMS over time. Quantification of relative LacZ splicing in untreated and MMS treated normalized to ‘no intron’ control, showing decrease in splicing activity within 30 minutes of MMS treatment. Mean± SEM, Student t-test, Asterisks show *p*-value (***) p<0.001. (B) Whole proteome analysis by mass spectrometry after MMS treatment. Pie chart of proteins quantified (4357 total, 3864 (89%) no change, 75 (2%) enriched, 418 (9%) depleted) (left) showing 9% (39) of the depleted proteins (green region of the zoomed red inset) are encoded from intron containing genes, which are predominantly ribosomal proteins (96%, 35) (Right pie chart). Depleted and enriched proteins are listed in **Supplementary Table S2**. (C) TOR signaling regulates Hsh155 foci formation. Treatment of cells with rapamycin (200nM, 2hr) reduces Hsh155 aggregate formation in MMS treated cells. Scale bar, 5μm. (D) Effect of temperature sensitive mutants of TORC1 subunit Kog1 and regulator Asa1 on Hsh155 relocalization at 25°C, 30°C. For C and D, three replicates, n>100, Mean± SEM, Fisher test p-value threshold **** p<0.0001.

Splicing flux in yeast is dominated by the production of ribosomal proteins (RPs), the majority of which encode an intron and whose transcripts account for ~90% of splicing reactions (Parenteau et al, 2011). It has long been known that RP genes are specifically repressed upon stress as part of a transcriptional program dubbed the Environmental Stress Response (ESR) (Gasch et al, 2000). Recent studies indicate that RP expression can also be regulated post-transcriptionally by selective splicing under stresses, including MMS (Gabunilas & Chanfreau, 2016; Parenteau et al, 2011). Our own whole proteome analysis revealed this bias to gene repression as 418 proteins were repressed and only 75 were more abundant after MMS treatment (**Fig. 7B; Supplementary Table S2**). Gene Ontology analysis of the cellular functions impacted highlights strong repression of transcription and translational processes (**Supplementary Table S3**). Specifically, our analysis confirmed a dramatic decrease in RP levels (**Fig. 7B; Supplementary Fig. S6C** and **Supplementary Table S2**); 9% of the down regulated proteins are encoded by intron-containing genes and this is driven by RP gene repression since all but four depleted and spliced proteins encode RPs (96%) (**Fig. 7B; Supplementary Fig. S6C**). Our results indicate that Hsh155 localizes to PQC sites precisely under conditions when its needed splicing function in the cell, RP production, is repressed. Indeed, our cell cycle analysis (**Fig. 6D**) showing foci in G1 but not G2/M is consistent with this model since ribosome production is required to progress through G1 (Bernstein & Baserga, 2004) while rRNA production at least is transiently decreased during mitosis (Clemente-Blanco et al, 2009). TOR signaling is known to regulate ribosome biogenesis normally and coordinate its repression under stress (Martin et al, 2004). Remarkably, co-treatment of cells with the TORC1 inhibitor rapamycin strongly suppressed MMS-induced Hsh155-GFP foci formation (**Fig. 7C**). Genetic perturbation of the TORC1 pathway through mutation of a Tor1/2 stabilizing chaperone *ASA1* (Stirling et al, 2011), or the essential TORC1 cofactor *KOG1* (Loewith et al, 2002), also significantly reduced the frequency of Hsh155-GFP foci (**Fig. 7D**). Overall, this suggested that RP repression, mediated by TORC1 signaling, could be influencing the dynamic behavior of Hsh155 in MMS.

### TORC1 influences sequestration of transcription regulators to PQC through Sfp1

The effects of TORC1 on RP gene expression are mediated through downstream effects on a group of transcriptional activators, including Hmo1, Ifh1, and Sfp1 (Reja et al, 2015; Schawalder et al, 2004; Xiao et al, 2011). Mutation of constitutive TORC1-regulated, RP gene transcriptional activators such as Hmo1 or Ifh1 either had no effect or caused a subtle decrease in the frequency of Hsh155-GFP foci (**Supplementary Fig. S6D**). Hmo1 and Ifh1 directly regulate RP gene expression but do not have an established role in the ESR. On the contrary, Sfp1 is a RP gene transcription factor that interacts with and is regulated by TORC1 signaling specifically under stress (Lempiainen et al, 2009). Normally, Sfp1 is displaced from RP gene promoters under stress and is localized to the cytosol to effect rapid adaptation of RP genes to stress (Jorgensen et al, 2004; Marion et al, 2004). In the absence of the Sfp1, RP genes are still transcribed but are not repressed under stress (Marion et al, 2004). To test the effects of removing this dynamic RP gene repression we assessed INQ formation in *sfp1∆* cells. Remarkably, while INQ protein aggregates marked by Apj1-GFP still formed normally in MMS-treated *sfp1∆* cells, Hsh155-GFP foci were completely abrogated and Hos2-GFP and Cmr1-GFP foci were also significantly reduced in frequency (**Fig. 8A and B**). Thus, loss of Sfp1 does not preclude the formation of aggregates; rather it controls the specific endogenous proteins that are recruited to the aggregates under stress.

**Figure 8.**
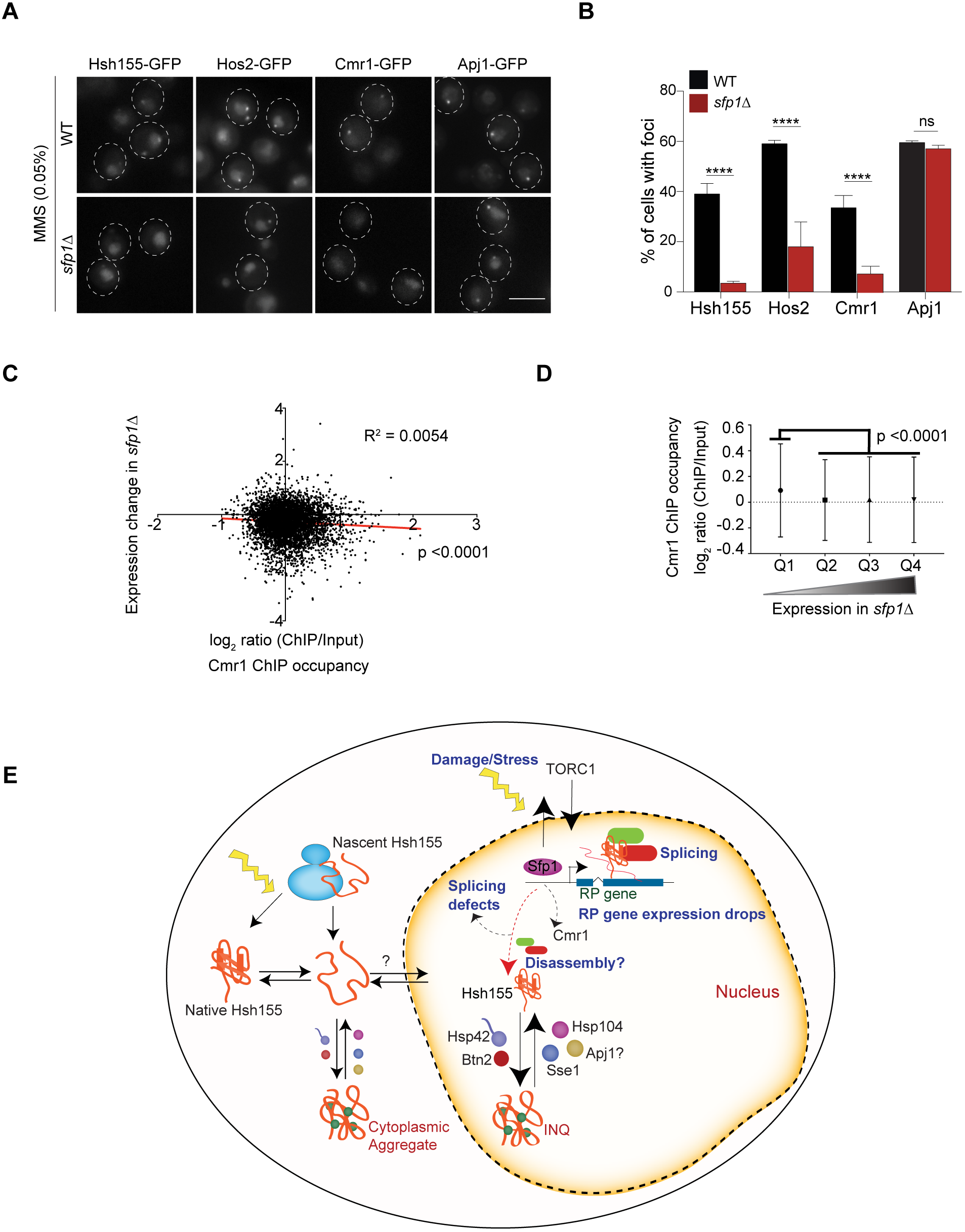
TORC1 influences sequestration of transcription regulators to PQC through Sfp1. (A) Sfp1 is required for Hsh155, Hos2, and Cmr1 relocalization but not for INQ formation. Representative images (left) showing effect of Sfp1 deletion on Hsh155, Hos2, Cmr1, and Apj1 localization. (B) Quantification of percentage of cells with Hsh155, Hos2, Cmr1, and Apj1 foci in WT (black bar) and *sfp1∆* (red bar). For B, three replicates, n>100, Mean± SEM, Fisher test p-value threshold **** p<0.0001. (C) Cmr1-occupied genes are significantly down regulated in *sfp1*∆. Scatter plot using microarray expression data in *sfp1∆* (Marion et al, 2004) and the Cmr1 ChIP occupancy data (All genes, n= 5549, listed in **Supplementary Table S4**) in wild type cells(Jones et al, 2016) and linear regression showing anticorrelation (marked by red line, R^2^= 0.0054; p< 0.0001) between expression changes in *sfp1∆* and Cmr1 occupancy in WT cells. (D) Quartiles were generated based on (C) presented as dot plots. The plots splits the *sfp1∆* expression changes into quartiles and shows that the lowest expression (Q1) in *sfp1∆* correlates with increased Cmr1 occupancy (ANOVA with Tukey’s test, *p*< 0.0001). Genes in each quartile listed in **Supplementary Table S4.** (E) Model illustrating stress-induced transcriptional changes at RP genes liberates transcriptional regulators such as Sfp1 regulated by TORC1 signaling, Cmr1, and spliceosomes leading to splicing defects. Subsequently, Hsh155 is sequestered in INQ and eventually cytoplasmic aggregates, which are regulated by chaperones until stress recovery (see main text).

Interestingly, like the spliceosome, Cmr1 and Hos2 have been linked to RP gene expression and bind to RP gene promoters (Jones et al, 2016). Together with our data, this suggests that a precipitous drop in RP gene expression might be involved in evicting Hsh155, Hos2, and Cmr1 from the chromatin and enabling their sequestration at PQC sites. To assess this possibility, we combined published microarray data on *sfp1∆* cells (Marion et al, 2004) with Cmr1 occupancy data in wild type cells by chromatin immunoprecipitation (ChIP) (Jones et al, 2016). This analysis showed that Cmr1-occupied genes are significantly down regulated in *sfp1∆* cells, supporting the idea that the two factors have common targets (**Fig. 8C** and **D; Supplementary Table S4**). Interestingly, analysis of the Cmr1 ChIP occupancy data also indicates that there is significantly more Cmr1 occupancy at spliced genes (average Cmr1 occupancy for 274 spliced genes −0.13 and average occupancy for all genes-0.046) in wild type cells (**Supplementary Fig. S6E** and **Supplementary Table S4).** Thus, genes regulated by Cmr1 and Sfp1 overlap and are enriched for spliced genes where Hsh155 will act. Together these data support a model where stress-induced transcriptional changes at RP genes and a precipitous drop in RP gene expression regulated by Sfp1 liberate spliceosomes and transcriptional regulators such as Hos2 and Cmr1. These factors can then be captured in protein aggregates in the nucleus, and eventually the cytoplasm, and sequestered until stress passes (**Fig. 8E**, see Discussion below).

## Discussion

### Hsh155 - a new INQ localizing protein

The INQ is a relatively poorly characterized PQC site for nuclear proteins. Our data establishes several new principles governing INQ formation and substrate protein recruitment. We show that INQ substrates can fall into at least two categories, those like Cmr1 that are wholly restricted to nuclear aggregates (Gallina et al, 2015), and those like Hsh155, Hos2 or Apj1 that, over time, accumulate in cytoplasmic foci. This could suggest that the capacity of INQ is regulated and proteins exceeding its ability to sequester are shunted to cytoplasmic aggregates sequentially making the latter an overflow compartment, or that both nuclear and cytoplasmic pool of proteins aggregate independently with different kinetics. A previous study (Miller et al, 2015) has indicated a simultaneous accumulation of substrates in both INQ and cytoplasmic PQC’s regulated by chaperones like Sis1 and nuclear import factors. Moreover, our tandem-fluorescent fusion data suggest that nascent Hsh155 accumulates in cytoplasmic aggregates compared with INQ and therefore we favor a model in which Hsh155 is not actively transported from INQ to cytoplasmic aggregates. In addition, since we show that INQ residents like Apj1 or Hos2 still localize to aggregates under conditions where Hsh155 does not (i.e. in MMS-treated *sfp1∆* cells for Apj1; spontaneously in *mec1∆tel1∆* cells for Hos2), there must be separate upstream signals for the formation of aggregates and the recruitment of specific endogenous proteins. Thus, the formation of the aggregate itself is insufficient to recruit a labile protein; rather signals which perturb Hsh155 structure or interactions must occur prior to its recruitment.

We also found that INQ markers Cmr1, Hos2, Apj1 and Pph21 affect the frequency of Hsh155 foci. However, based on the annotated functions of these proteins we propose that there may be different mechanisms by which this occurs. Apj1 is homologous to Hsp40, a molecular chaperone, and we predict that it plays a role in stabilizing soluble Hsh155 in the nucleus and cytoplasm. Cmr1 and Hos2 are now known to affect transcription and ribosomal protein gene expression (Jones et al, 2016) and thus deletions would disrupt the levels of spliced transcripts, potentially sensitizing cells to sequester Hsh155 in PQC’s under stress. Finally, Pph21 opposes TORC1 dependant phosphorylation of Tap42 (Jiang & Broach, 1999), which would be partly alleviated in *pph21∆* cells and associated with a stronger TORC1 signal to Sfp1 under stress. This potentially explains why rapamycin treatment blocked Hsh155 foci formation, while *pph21∆* cells were sensitized to accumulate Hsh155 foci. Thus, while each of these INQ proteins fits into a model of stress-signaling induced transcriptional changes leading to protein aggregation, there are many remaining questions about how and why this subset of proteins are sequestered at the INQ and whether they are inactive aggregated substrates or are exerting their enzymatic activities (i.e. lysine deacetylation by Hos2, or S/T dephosphorylation by Pph21) within the INQ.

The role of molecular chaperones at protein aggregates is clearer and we identify well known disaggregase and aggregases affecting Hsh155 deposition at aggregates. Interestingly, as with Cmr1, Hsh155 localization is regulated by both Btn2 and Hsp42. Unlike Cmr1, Hsh155 is also deposited at cytoplasmic aggregates and this behavior is also abolished in cells lacking *BTN2* or *HSP42.* This supports a common function for Btn2 and Hsp42 in aggregation processes, although whether they are functionally redundant, work in a complex, or act through their influence on another protein is unknown. Deletion of the disaggregase *HSP104* dramatically increased the frequency of Hsh155 foci in the presence of MMS indicating possible role in refolding and reactivation of Hsh155 after stress recovery. *SSE1* deletion led to spontaneous Hsh155 foci suggesting a constitutive role for Sse1 in spliceosomal integrity.

### Linking transcriptome remodeling and protein quality control

The coordination of changes in the transcriptome and proteome are essential to reestablish cellular homeostasis after stress. Temporary protein sequestration or turnover in aggregate structures is one way that such homeostasis is achieved (Saarikangas & Barral, 2016; Wallace et al, 2015). The degree to which the DNA damage response to MMS treatment affects these peripheral stress responses is unknown. Here we find that neither entry into S-phase nor signaling through Tel1 or Mec1 are required to form Hsh155 INQ foci, suggesting that Hsh155 aggregation is largely independent of canonical DNA damage signaling emanating from the recognition of MMS-damaged DNA.

Instead, we propose that the transcriptional response to MMS is the ultimate driver of Hsh155 aggregation. Spliced transcripts are dramatically affected by MMS treatment because RP production is shut-down and RP genes encode the majority of spliced transcripts in yeast. Thus, Hsh155 is no longer engaged in bulk splicing at RP genes under stress. This situation may also apply to Cmr1 and Hos2, which bind to RP genes during transcription and may be evicted during transcriptional repression (Jones et al, 2016). These dynamics at RP genes are initiated by TORC1-dependant relocalization of Sfp1 (Jorgensen et al, 2004; Marion et al, 2004) and we show that either TORC1 inhibition by rapamycin or *kog1-ts, asa1-ts,* and *SFP1* deletion strongly repress Hsh155 or Hos2 or Cmr1 aggregation in MMS. We also show complete abrogation of Hsh155 foci formation by inhibiting new translation using CHX with MMS. Since it is known that CHX can also affect ribosome biogenesis (Reiter et al, 2011), we suspect that the repression of RP synthesis by CHX along with whole protein synthesis shut off, might have significantly contributed to Hsh155 aggregate suppression under stress. Importantly, INQ protein aggregates still form when Sfp1 is absent, as marked by Apj1-GFP. Thus, only specific substrates of INQ are impacted by changing the transcriptional dynamics of the stress response. This is important since it may suggest that Hsh155 and Hos2 are not simply aggregation prone in MMS but instead move to INQ, or not, based on the transcriptional needs of the cell.

Previous groups have recognized that INQ structures localize adjacent to the nucleolus (Miller et al, 2015; Tkach et al, 2012) and our data directly links the constituents of INQ to ribosome production (i.e. through the cell cycle, Sfp1, TORC1). While the peri-nucleolar localization of INQ may therefore be significant, the relationship of these structures to ribosome assembly is currently unknown. Recent identification of quality control mechanisms for unassembled ribosome subunits (Sung et al, 2016) and that ribosomal stalling of nascent proteins induces selective aggregation (Yonashiro et al, 2016) raises the hope that this can be explored in the near future. Another outstanding question relates to the signal that causes Hsh155 to localize to aggregates while its interacting partners in the spliceosome do not. Whether this is an inherent chemically sensitive property of Hsh155 protein or it is a regulated disassembly reaction aimed at suppressing splicing is currently unknown. Our study highlights INQ as an immediate repository for factors perturbed by RP gene repression and, by linking dynamic transcriptional changes to PQC, raises important questions about how cells coordinate the assembly and disassembly of chromatin-associated protein complexes during stress and recovery.

## Materials and Methods

### Yeast growth, manipulation and analysis

Yeast strains were grown in standard rich media YPD or synthetic complete (SC) medium unless otherwise indicated. Serial dilution assays were performed as described (Stirling et al, 2011). Standard MMS treatments (unless indicated), were at 0.05% for 2hr (~99% Sigma). All other chemical concentrations and treatment times are indicated in each figure or legend. **Supplementary Table S5** contains a list of yeast strains, primers and plasmids used.

### Live cell imaging and CIN-GFP screen

Genes with reported genome instability (Stirling et al, 2011; Stirling et al, 2014) were obtained as GFP fusions (Huh et al, 2003). Actively growing cells were exposed to H_2_O_2_ (2mM) or MMS (0.05%) for 2 hours in batches of 12 strains in well plates before mounting on concanavalin A (ConA) treated (Stirling et al, 2012), Teflon masked 12-well slides. For UV exposure, untreated cells were mounted in 12-well slides and the droplets were irradiated (500 J/m^2^) in a stratalinker. Irradiated slides were stored in a humid chamber until imaging. The imaging screen was conducted on a Zeiss axioscop at 100x magnification and candidate relocalizations were retested in triplicate. Imaging of the treated strains after the screen was performed live on a Leica DMi8 microscope at 100x magnification using ConA treated slides. VHL-mCherry aggregate induction was done as described (Miller et al, 2015).

### Cell cycle anaylsis

Logarithmic cultures of GFP tagged Hsh155 cells were grown at 30°C and treated with α-factor (1mg/ml, 1hr) for G1/S arrest and with nocodazole (15μg/ml, 1hr) for G2/M arrest before MMS treatment (0.05%, 2hr). Arrested and MMS treated cells were then imaged as described above.

### Image analysis and statistical methods

The images were acquired on Leica DMi8 microscope using MetaMorph Premier acquisition software and post processed using ImageJ. For all microscopy experiments, the significance of the differences was determined using Prism5 (Graphpad) or R. For intensity measurements samples were compared with t-tests or ANOVA; graphpad performs F-tests for variance as part of this analysis. For comparisons of proportions, Fisher tests were used and p-values were Holm-Bonferroni corrected in the event of multiple comparisons. Sample sizes were determined *post hoc* and are listed in the figure legends.

### FRAP analysis

FRAP experiments were done using an Olympus FV1000 confocal imaging exactly as described (Chao et al, 2014). Hsh155 and Hos2-GFP tagged cells were grown to log-phase and treated with MMS (0.05%) for 2hrs. Cells with foci in the nucleus were selected for imaging. FRAP images were collected on an Olympus FV1000 microscope with Olympus Fluoview version 3.0. Hsh155 and Hos2-GFP INQ foci were bleached, and the recovery of fluorescence in the bleached region of interest (ROI) was monitored every 5 seconds (s). The bleaching experiment was performed using a 488-nm laser using 40% bleach laser power and 1 frame bleach time. All fluorescence normalization was automated using R and R Studio3 (RStudio Team (2015). RStudio: Integrated Development for R. RStudio, Inc., Boston, MA URL http://www.rstudio.com/). Background fluorescence was monitored in three ROIs, averaged, and removed from the bleached ROI. Three control ROIs in the nuclei of neighbouring cells were monitored, averaged, and normalized to the prebleach ROI (T_0_) and the fluorescence loss over time was calculated and added back to the bleached ROI. Normalized bleached ROI fluorescence data was transformed by setting the prebleached ROI to 100 %, and the post-bleached ROI to 0 % to allow for all FRAP curves to be combined. Data was fit using Prism6 (GraphPad) by one phase association nonlinear regression. Mobile fraction and t_1/2_ values were calculated and obtained from the values output by Prism6.

### Western blotting

For western blots, whole cell extracts were prepared by tricholoroacetic acid extraction and blotted with anti-GFP (ThermoFisher) or anti-PGK1 (Abcam) essentially as described (Gallina et al, 2015).

### Splicing efficiency assay

Splicing assay protocol was adapted and performed as previously described (Galy et al, 2004). All measurements were taken with individual transformants in triplicate. Cells were struck as a patch on SC medium without leucine, then replica plated to glycerol-lactate-containing synthetic complete medium without leucine (GGL-leu). Cells from each patch were inoculated in liquid GGL-leu media for 2 hours at 30°, then induced with final 2% galactose for 1.5 hours before treatment with final 0.05% methyl methanesulfonate (MMS). Time points were taken at 30, 60, and 120 minutes post-MMS treatment. Cells carrying reporters were lysed and assayed for β-galactosidase assay using Gal-Screen^™^ β-Galactosidase Reporter Gene Assay System for Yeast or Mammalian Cells (Applied Biosystems) as per manufacturer’s instructions, and read with a SpectraMax i3 (Molecular Devices). Relative light units were normalized to cell concentration as estimated by measuring OD600.

### Whole proteome analysis by mass spectrometry (MS)

Logarithmic cultures of BY4741 wildtype (WT) strain grown at 30°C with or without MMS (0.05% for 2hr) treatment were pelleted and frozen. Frozen pellets were lysed, reduced, alkylated, trypsin digested, and purified using the SP3 method (Hughes et al, 2014) with modifications (Hughes et al, 2016). Samples were analyzed as detailed in (Hughes et al, 2016); briefly, prepared peptide samples were labeled with individual tandem mass tags (TMT, Pierce), combined in sets of 10, and subjected to offline high pH fractionation/concatenation, then fractions (12) were analyzed by reverse phase nano-electrospray liquid chromatography on a Orbitrap Fusion Tribrid MS platform (Thermo Scientific) using MS3 scanning. Mass Spectrometry Data Analysis: Data from the Orbitrap Fusion were processed using Proteome Discoverer Software (ver. 2.1.0.62). MS2 spectra were searched using Sequest HT against the UniProt *Saccharomyces cerevisiae* proteome database appended to a list of common contaminants (6,752 total sequences). Data were filtered at the peptide spectral match-level to control for false discoveries using a q-value cut off of 0.05 as determined by Percolator. This less-stringent filter was applied to maximize sensitivity, relying on the statistical analyses during peptide quantification to further control for the potential generation of false conclusions within the final data set. As a result, the final quantitative set of hits that displays significant variance between sample types is enriched in multi-peptide identified, high confidence proteins. A total of 4357 proteins were reproducibly quantified, the proteins with significant depletion or enrichment are list in **Supplementary Table S2**.

Bioinformatic and Statistical Analyses: Data sets generated in Proteome Discoverer were exported and analyzed with a combination of scripts built in R designed in-house. Contaminant and decoy proteins were removed from all data sets prior to analysis. Unless stated otherwise, quantification was performed at the peptide level as discussed previously(Suomi et al, 2015). Data Availability: The mass spectrometry proteomics data have been deposited to the ProteomeXchange Consortium via the PRIDE partner repository (Vizcaino et al, 2016; Vizcaino et al, 2014) with the dataset identifier PXD004459 (Username, Password: Udzf7jqI).

## Acknowledgements

We acknowledge Grant Brown, Shay Ben-Aroya, Philip Hieter, Benoit Palancade, Daniel Kaganovich for yeast strains and plasmids, Chhabi Govind for help accessing the Cmr1 ChIP data and members of the Stirling lab for helpful discussions. P.C.S. is a Canadian Institutes of Health Research (CIHR) New Investigator, and a Michael Smith Foundation for Health Research Scholar. C.S.H and G.B.M. acknowledge support from the British Columbia Cancer Foundation. This work is supported by operating grants from CIHR (MOP-136982) and the Natural Sciences and Engineering Research Council of Canada to P.C.S (RGPIN 2014-04490).

## Author contributions

V.M. and P.C.S designed the project and wrote the manuscript. V.M., A.S.T., A.K.H., C.S.H., and P.C.S. generated the data. V.M., A.S.T., K.L.M., A.K.H., C.S.H., G.B.M., C.J.R.L., and P.C.S. analyzed the data.

## Supporting Information

**Fig S1.**
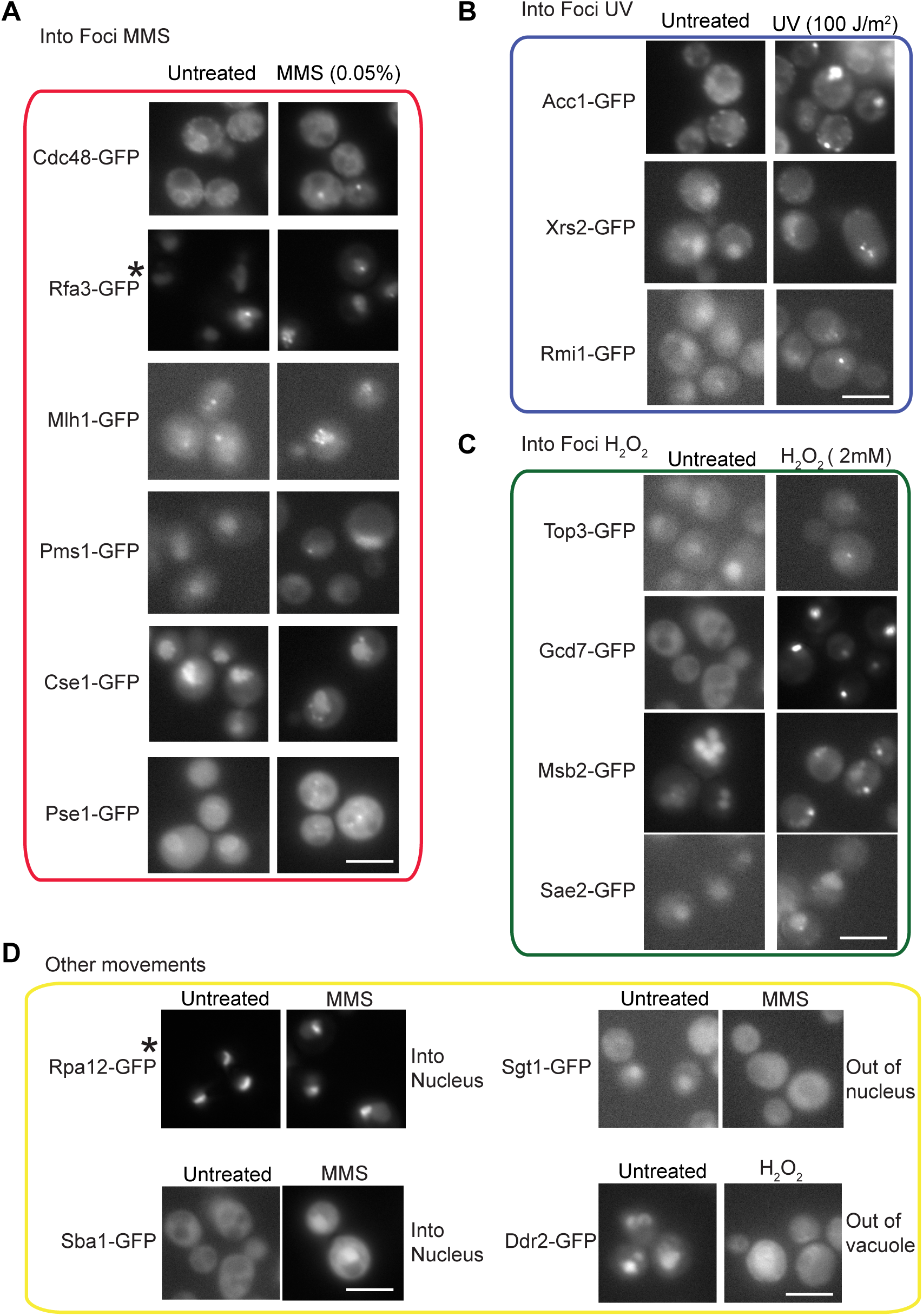
Novel protein relocalization behaviors in the CIN mini array screen. Movement types are grouped as indicated, (A) MMS induced foci (red box), (B) UV induced foci (blue box), and (C) H2O2 induced foci (green box). (D) Non-foci movements as shown (yellow box). Only those not reported in Tkach et al., 2012 and a single condition is shown for clarity, however proteins may move in response to multiple stresses – for a complete list, see **Supplementary Table S1**. In each case, the untreated localization is the left panel, and the indicated treatment is shown at right. Hsh155-GFP relocalization is not shown (see **Fig 1**). The asterisks indicate that this is a representative of multiple members of a protein complex. In the case of Rfa3-GFP, its partners Rfa2 and Rfa1 also showed more foci in MMS (see **Table S1**). In the case of Rpa12-GFP, its partners Rpa14 and Rpa34 also showed observed increases in nucleoplasm versus nucleolar fluorescence in MMS. The Rpa12-GFP observation was confirmed by measuring nucleoplasmic fluorescence by colocalizing with HTA2-mCherry and quantifying the GFP signal in the non-nucleolar nuclear area (n = 17, t-test p = 0.016). Scale Bar, 5μm.

**Fig. S2.**
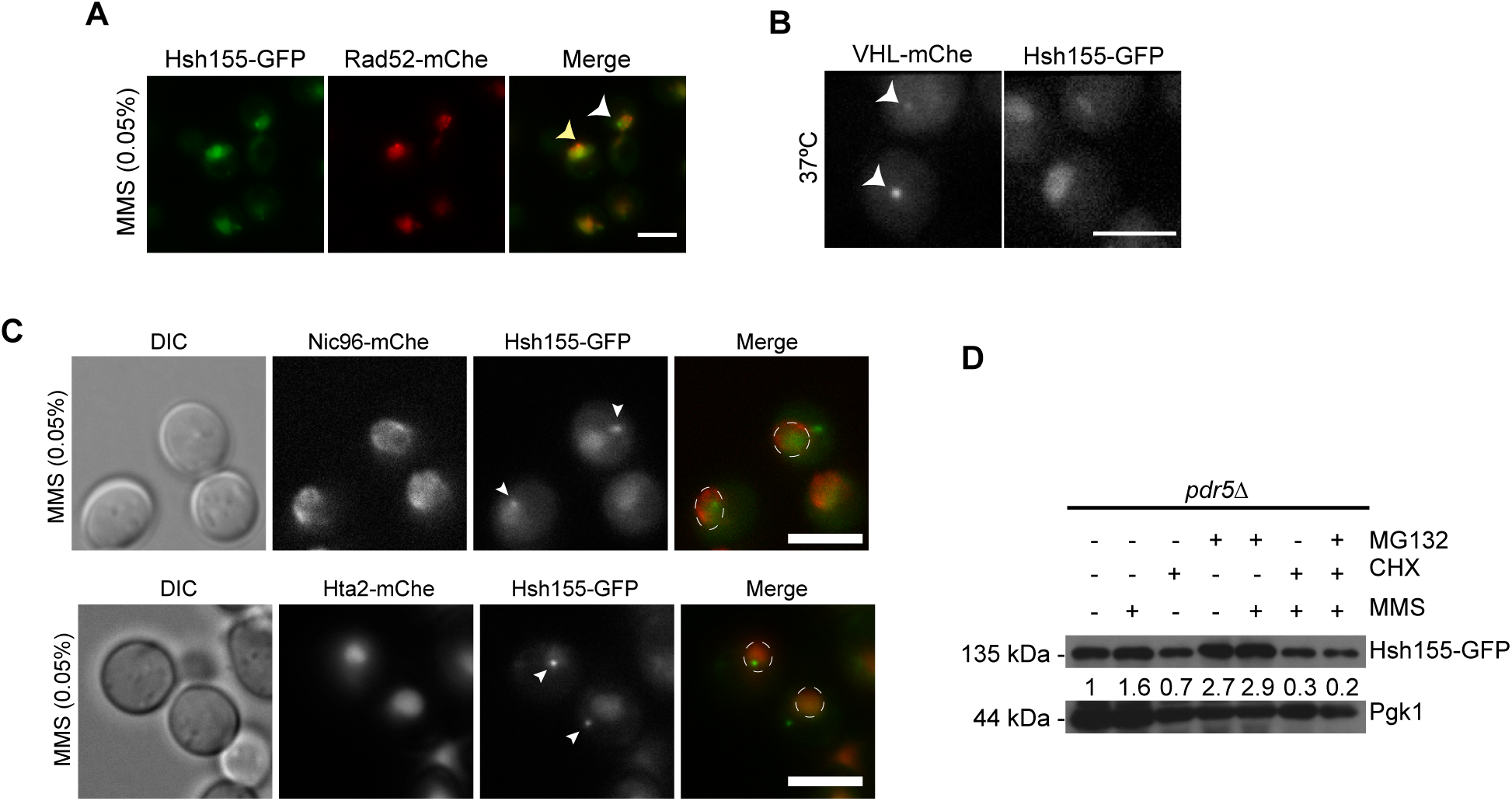
Characterization of Hsh155 relocalization. (A) Hsh155-GFP foci (white arrowhead) do not colocalize with Rad52-mCherry foci (yellow arrowhead) after 2hr of MMS treatment. (B) Hsh155 do not form foci with heat stress at 37°C, while a VHL control does (arrowheads). (C) Localization (white arrowheads in merge) of Hsh155-GFP in nucleus and cytoplasm, nuclear periphery marked with Nic96-mCherry (mChe) (top) and chromatin marked with Hta2-mCherry (bottom) in MMS. Bar, 5μm. (D) Hsh155 protein levels relative to Pgk1 levels (as indicated) in cells treated with MMS (0.05%) and/or CHX (200μg/ml), MG132 (80μΜ) alone or together for 2hrs. Hsh155 protein levels appear to be comparable to untreated in all treatments.

**Fig. S3.**
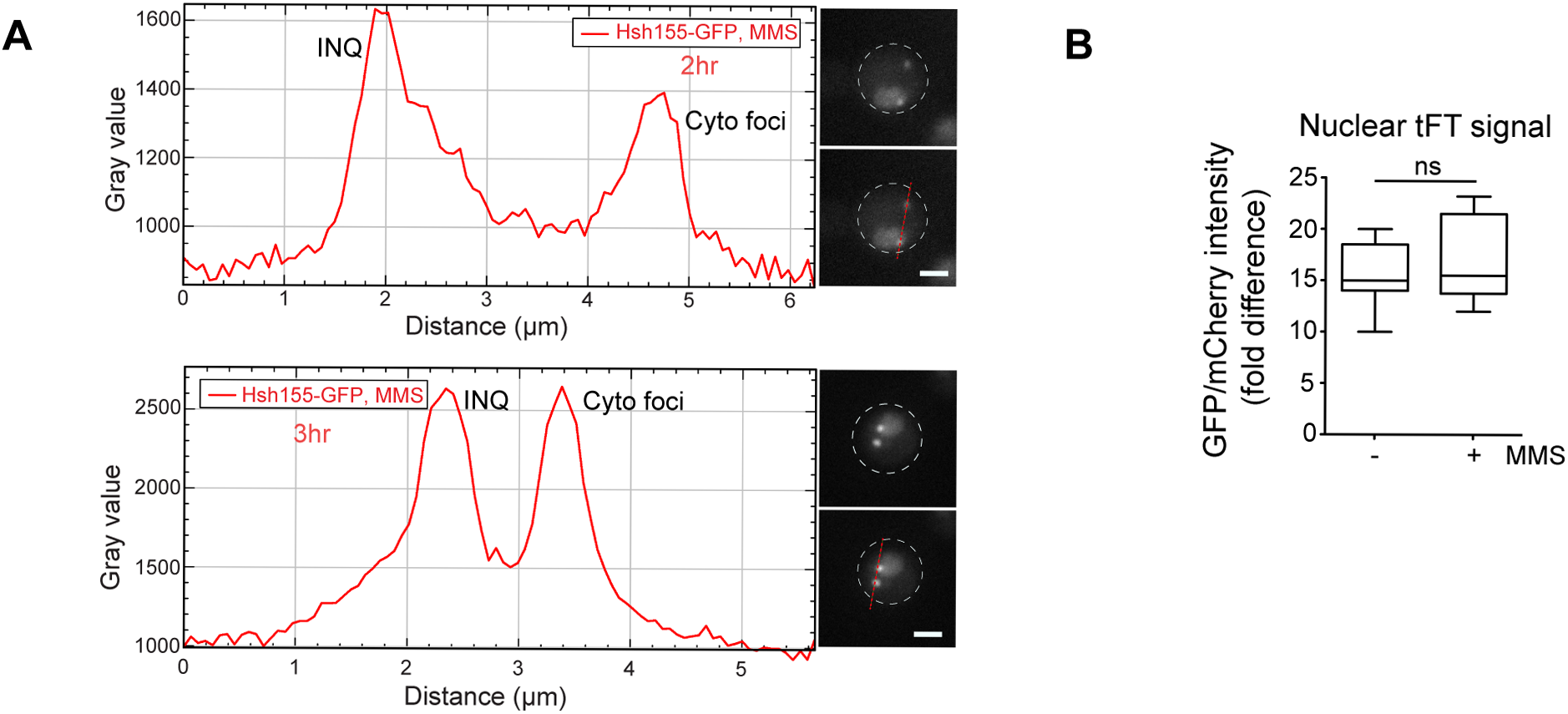
Dynamics of Hsh155 relocalization under stress. (A) Line scan plots of Hsh155-GFP intensities in INQ and cytoplasmic foci (depicted by dotted red line in representative images) at two (above) and three (below) hours of MMS treatment. Bar, 2μm. Representative graphs shown in **Fig. 3C.** (B) Quantification of fold differences in Hsh155 tandem fusion to a fluorescent timer shows non-significant (ns) differences in lifetime between untreated and MMS treated nucleoplasm.

**Fig. S4.**
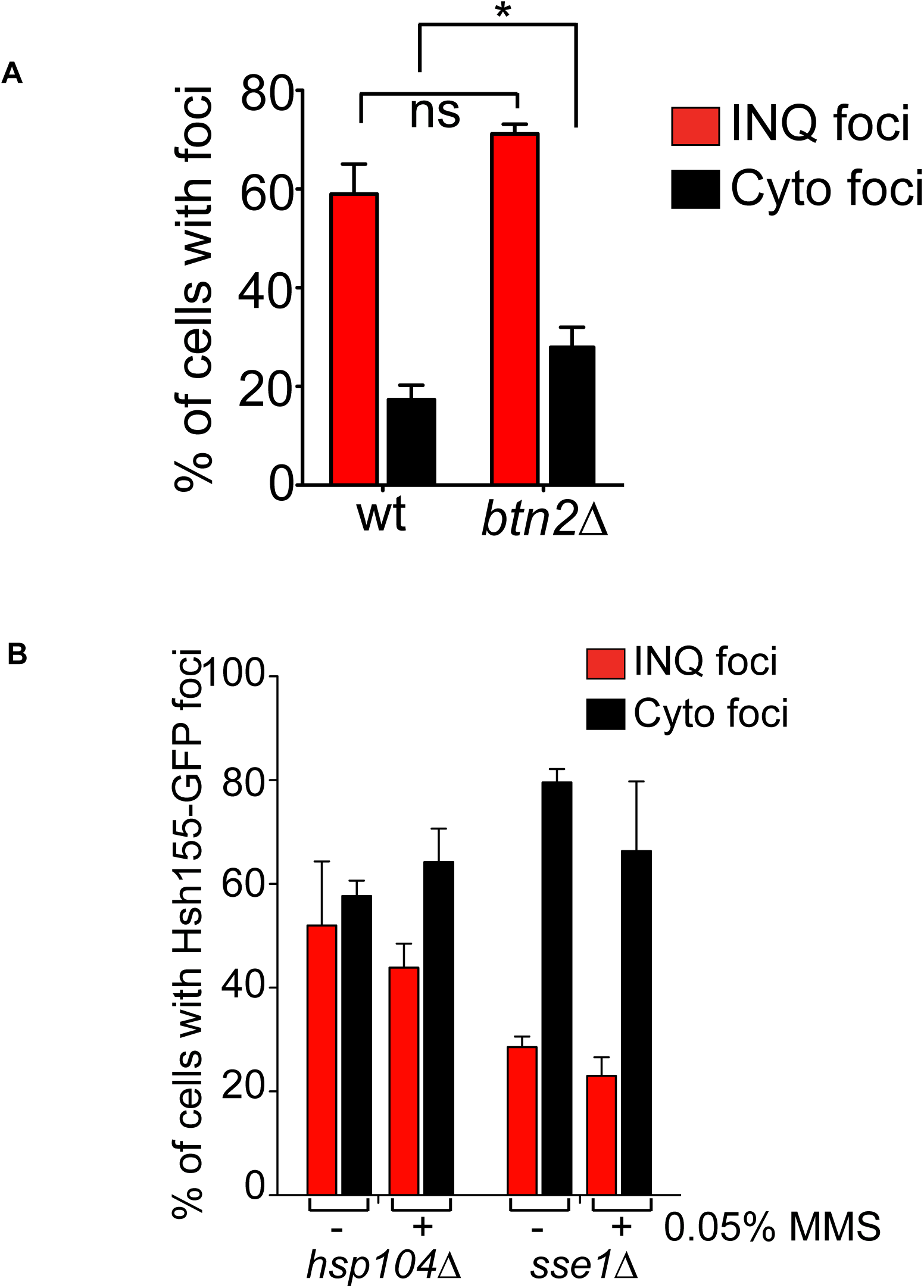
INQ and Cytoplasmic site foci distribution of Hsh155 in chaperone mutants. (A) Effect of deleting *BTN2* on INQ and cytoplasmic foci distribution. No significant difference in both nuclear and cytoplasmic foci distribution in *btn2* mutants compared to WT. Shown are the mean values from three independent experiments ± SEM with at least 100 cells each. Fisher test *p*-value thresholds, ns = non-significant and * p<0.05. (B) Effect of deleting *HSP104, SSE1* on INQ and cytoplasmic foci distribution. Percentage of cells with cytoplasmic foci (black bar) increased in *hsp104* mutants in MMS and in both untreated and MMS treated cells in *sse1* mutants. Mean values from three independent experiments ± SEM with at least 100 cells each.

**Fig. S5.**
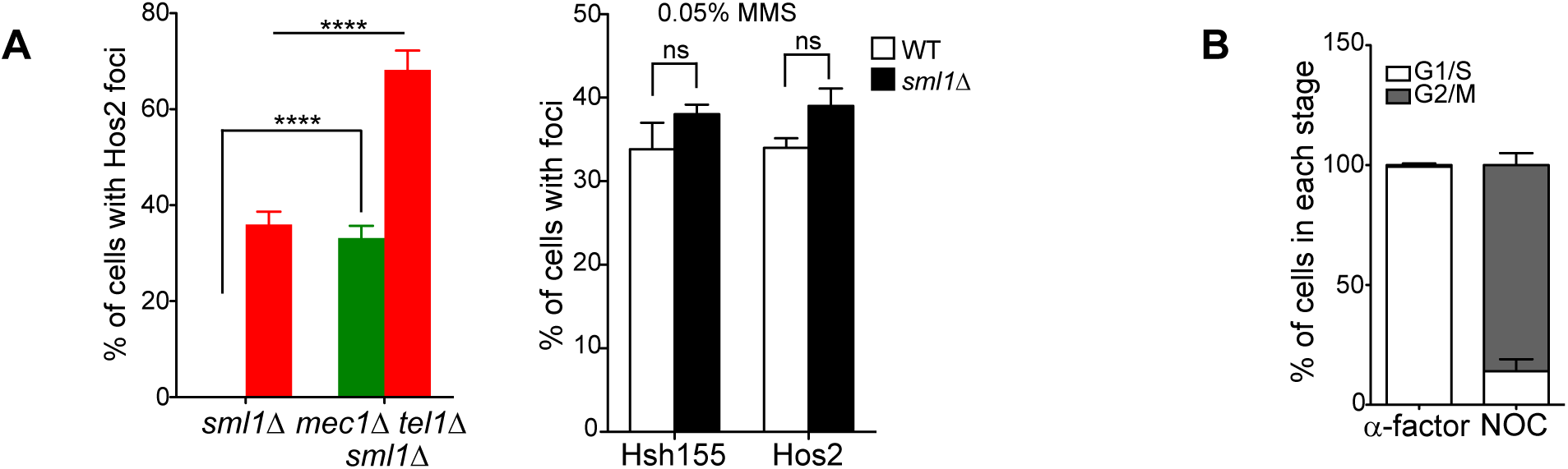
Cell cycle dependency of Hsh155 relocalization. (A) Loss of *MEC1* and *TEL1* influences Hos2 aggregate formation. Bar graph (left) shows significant difference in Hos2 localization in *mec1∆tel1∆.* in both untreated (green bar) and MMS treated (red bar). Representative image shown in **Fig. 6C.** Bar graph (right) showing no significant difference in localization of either Hsh155 or Hos2 in *sml1∆* control for *mec1∆tel1∆sml1∆* mutants. All quantifications: three replicates, n> 100, Mean± SEM, Fisher test, Asterisks show *p*-value thresholds, ns = non-significant; **** p<0.0001. (B) Percentage of cells arrested in G1/S (white) and G2/M (grey) in α-factor and nocodazole (NOC) treated cells as judged by budding index. Three replicates, n> 100.

**Fig. S6.**
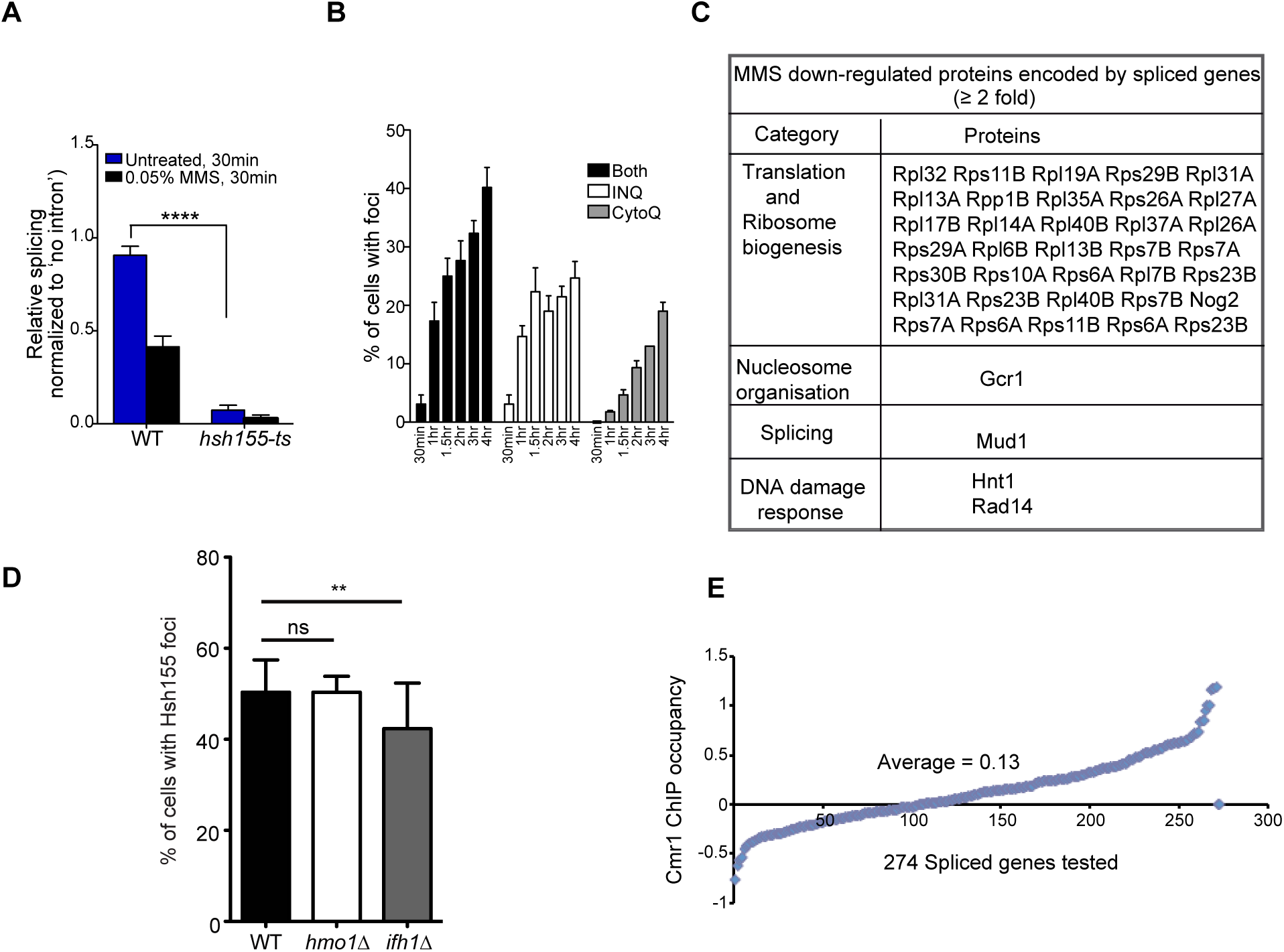
Transcriptional remodeling under stress linking to PQC site targets. (A) Splicing efficiency in *hsh155-ts* (used as a positive control) at 30°C compared to wild type is significantly reduced with and without MMS treatment. Quantification of relative splicing in untreated (blue bar) and MMS treated (black bar) normalized to ‘no intron’ control. Results were compared with a t-test, ****p<0.0001. (B) Hsh155 protein aggregates at PQC sites are not formed until one hour of MMS treatment. Percentage of cells with foci in INQ (white bar), cytoplasm (grey bar) or both (black bar) after MMS treatment at different times as indicated. (C) List of MMS induced down regulated proteins encoded by intron containing genes in the whole proteome enrichment analysis by mass spectrometry, which majorly includes all ribosomal proteins. (D) Effect of RP gene transcription regulators Hmo1 and Ifh1 on Hsh155 relocalization after MMS treatment. Shown is the mean± SEM of three replicates, n>100, Fisher test p-value thresholds, ns= non-significant, **p<0.01. (E) Scatter plot showing distribution of 274 spliced genes (X axis) with their corresponding Cmr1 ChIP occupancy (Y axis-log_2_ ratio ChIP/Input). The plot indicates a highly significant and larger Cmr1 occupancy in WT cells (Jones et al, 2016) at spliced genes. Average Cmr1 occupancy for 274 spliced genes −0.13 and average occupancy for all genes- 0.046. Two-tailed Student t-test, p<0.0001 (All the genes n= 5549, and spliced genes n=274 compared listed in **Supplementary Table S4**).

**Table S1.** List of GFP tagged CIN gene mini array screen proteins showing localization changes after MMS, H2O2 or UV treatment.

**Table S2.** Whole proteome abundance data after MMS treatment.

**Table S3.** Gene Ontology enrichment for depleted proteins upon MMS treatment

**Table S4.** Comparison of the Cmr1 ChIP occupancy data in WT cells (Jones et al, 2016) and complete microarray expression data in *sfp1∆* strain (Marion et al, 2004).

**Table S5.** Yeast strains used in this study

